# Subnet Communicability: Diffusive Communication Across the Brain Through a Backbone Subnetwork

**DOI:** 10.1101/2023.09.20.558638

**Authors:** Jonathan Parlett, Abhishek Jeyapratap, Ali Shokoufandeh, Birkan Tunc, Yusuf Osmanlioglu

## Abstract

One of the fundamental challenges in modern neuroscience is understanding the interplay between the brain’s functional activity and its underlying structural pathways. To address this question, we propose a novel communication pattern called *subnet communicability*, which models diffusive communication between pairs of regions through a small, intermediary subnetwork of brain regions as opposed to spreading messages through the entire network. We demonstrate that subnet communicability strengthens coupling between the structural and functional connectomes better than previous models, including communicability. Over two large datasets, we show that the optimal subnetwork is consistent across the population. Subnet communicability provides new insights into structure-function coupling in the brain and offers a balance between redundancy in message passing and economy of brain wiring.

## 1 Introduction

The human brain is a complex network of interconnected neural elements that can be considered as an information processing network. At the macroscale, the human connectome maps the connectivity between brain areas [6]. Using magnetic resonance imaging (MRI), the connectivity between brain regions can be described in terms of structural relationships between gray matter regions that denote anatomical connectivity through white matter pathways [23], or functional relationships capturing statistical patterns of co-activation over time that correspond to communication between these regions [3]. Functional connectivity and rich network dynamics are influenced and constrained by anatomical connections and brain network topology [4, 12]. Understanding the dynamics of functional interactions between brain regions with no direct anatomical connections [13] is an open challenge in modern neuroscience.

Several network communication models have been devised to explain the relationship between observed functional connectivity (FC) and underlying structural connectivity (SC) in the brain. By modeling the possible neural communication pathways shaped by structural connections, these models aim to describe the polysynaptic interactions between anatomically unconnected brain regions [1]. Using a particular candidate model, each individual’s structural connectivity matrix can be augmented into a communication matrix, i.e., simulated functional connectivity, that represents the connectivity between all pairs of regions in the brain [21]. This simulated functional connectome is then compared with the empirical functional connectome to quantify similarity.

One of the earliest models proposed was shortest-path, where indirect communication between region pairs occurs through a minimum number of intermediate hub regions [12]. When considered on a weighted connectome, a variation of shortest-path was suggested where messages travel through the path with strongest connectivity [24]. These were considered de facto communication models in the brain as the idea of communication through a single optimal path is in alignment with the established wiring economy of the brain [2]. However, the efficacy of shortest path was questioned since its calculation requires having a complete knowledge of the network topology, which can be considered implausible for local neural elements in the brain [22]. Additionally, since the model assumes communication happens through a single pathway, it lacks the redundancy that is necessary for robust communication.

To overcome these limitations, communication patterns such as path transitivity and search information were proposed. These models advocate communication through parallel pathways that detour around the shortest path [11] and demonstrate better structure-function coupling than the shortest-path model. A decentralized communication pattern called communicability [7, 9] models communication occurring diffusively through all possible pathways in parallel. Communicability was recently shown to increase coupling between the structural and functional connectome better than other communication patterns including shortest-path, search information, and path transitivity [17, 20, 28]. Despite its success in better explaining structure-function coupling and accounting for redundancy, communicability contradicts the established economy of brain wiring as it does not restrict volume of information sent when flooding the entire network for communication.

In this study, we propose a novel network communication model called *subnet communicability*. Our proposed model aims to limit the redundancy of communicability and provide an efficient wiring economy while still offering a decentralized communication scheme. We achieve this by restricting diffusive communication to occur through a backbone subnetwork of a considerably smaller size, which is connected to the rest of the network. We systematically evaluate subnetworks of varying sizes and investigate the set of regions that constitute subnetworks with the highest structure-function coupling across individuals. We also analyze the contribution of functional systems to these subnetworks. Our proposed model offers insight into the underlying mechanisms used to process information in the brain and its biological neural signaling patterns.

## 2 Methods

### 2.1 Dataset and Preprocessing

We evaluated our methods on 200 unrelated, healthy young adults (96 males) in the age range [22,35] from the S1200 Young Adult Open Access dataset of the Human Connectome Project (HCP) [25]. To test the generalizability of our approach, we repeated our experiments on 261 healthy individuals (140 males) in the age range [22,86] from the 1000Brains dataset [5]. Structural and functional connectomes used in our analysis were provided open source in [8, 15], which were derived from diffusion-weighted MRI (dMRI) and resting-state functional MRI (fMRI) data. Connectomes were generated using the Schaefer atlas with 100 regions [18], where structural connectivity was obtained through probabilistic tracking with 10M streamlines and functional connectivity was obtained by calculating Pearson’s correlation over the BOLD signal. The reader is referred to [8] and [15] for more details of the data processing pipeline.

### 2.2 Overview of Structure-Function Coupling

Taking the structural connectivity of a subject as the basis, we calculated simulated functional connectivity by using a communication model. We then calculated Pearson’s correlation between the resulting simulated connectome and positive empirical functional connectome to quantify structure-function coupling (SFC). Higher correlation indicates better coupling (Fig. 1, left).

**Fig. 1.**
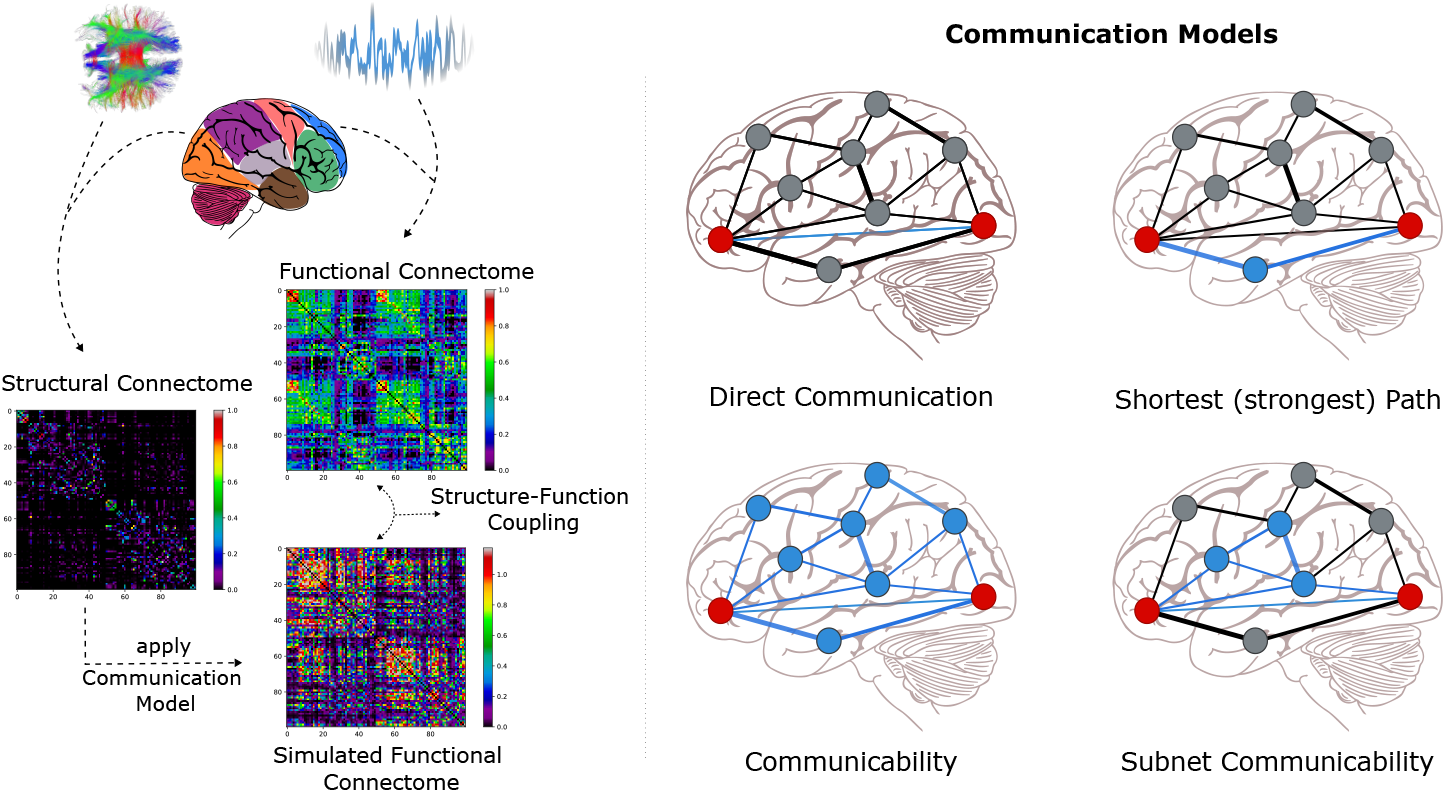
Overview of the method: (left) Using the structural connectome of an individual, one of the communication models is applied to obtain a simulated functional connectome, which is then correlated with the positive functional connectome to calculate structure-function coupling. **(right)** Visual representation of explored communication models. Red nodes denote source and destination regions in the brain network. Blue edges represent the pathway that the message travels through, and edge thickness accounts for the strength of the structural connection.

### 2.3 Communication Models

In our structure-function coupling analysis, we propose subnet communicability and compare it to three other communication models (Fig. 1, right). Each communication model is applied on a subject’s SC matrix *W*, where *W*_*ij*_ is the strength of structural connections between region pairs.

#### Communicability

Communicability, which is the basis of our proposed model, utilizes a broadcasting approach where signals are simultaneously propagated through all possible regions in the network [9]. Unweighted communicability between nodes *i, j* is computed by calculating the number of walks between them scaled relative to path length *k* as 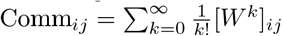. To account for the influence of connection strength in weighted SC matrices, weighted communicability is computed over the normalized *W* ^*′*^, where 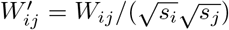, and *si* is the strength of node *i*.

#### Subnet Communicability

Extending the definition of weighted communicability, which propagates a signal across all possible fronts, we propose subnet communicability which propagates a signal only through a subset of regions that constitute a backbone network. Given a graph *G* = (*V, E*) and a subset of nodes *H ⊆ V* to constitute a subnetwork, we compute subnet communicability between two regions *i, j ∈ V* by first forming the subgraph composed of the nodes *H ∪ {i, j}* and their associated edges in *E* with corresponding SC matrix *W*_*H*_ . We then calculate connectivity by 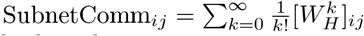. Repeating this process for all node pairs yields the subnet communicability matrix.

#### Shortest Path

The shortest path model routes information deterministically using a centralized strategy [21]. Given a weighted *W*, connectivity between *i* and *j* is given by *P*_*ij*_ = *e*_*iu*_ + … + *e*_*vj*_, the sum of edge weights in the strongest path between *i* and *j*, it is computed using an all pairs shortest path algorithm given *W* ^*′*^ as input, where 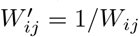

#### Direct Communication

Used as a baseline for comparing the efficacy of the other communication models, this model accounts for communication happening only between regions that are anatomically connected to each other.

## 3 Results

### SFC analysis on subnetwork size

We first explored the SFC for subnet communicability using varying network sizes ranging from 1 to 75 regions across a subset of the HCP dataset consisting of 40 individuals. We randomly sampled nodes to constitute subnetworks 1000 times at each size (except for size 1 where the number of possible subnetworks is 100), while ensuring that each node in the graph has a connection to at least one of the nodes in the resulting subnetwork. Evaluating the highest SFC scores for subjects across random samplings at each size, we observed that SFC achieves a peak for subnetworks of size 3 with a mean score of *r* = 0.35 across the subjects and steadily decays with increasing subnetwork sizes, converging to *r* = 0.27 for standard communicability (Fig. 2, left). For a single subject, SFC scores of all randomly sampled subnetworks of varying sizes are also shown (Fig.2, right). Initial results demonstrate that higher SFC can be achieved using subnet communicability relative to standard weighted communicability that uses the entire network.

**Fig. 2.**
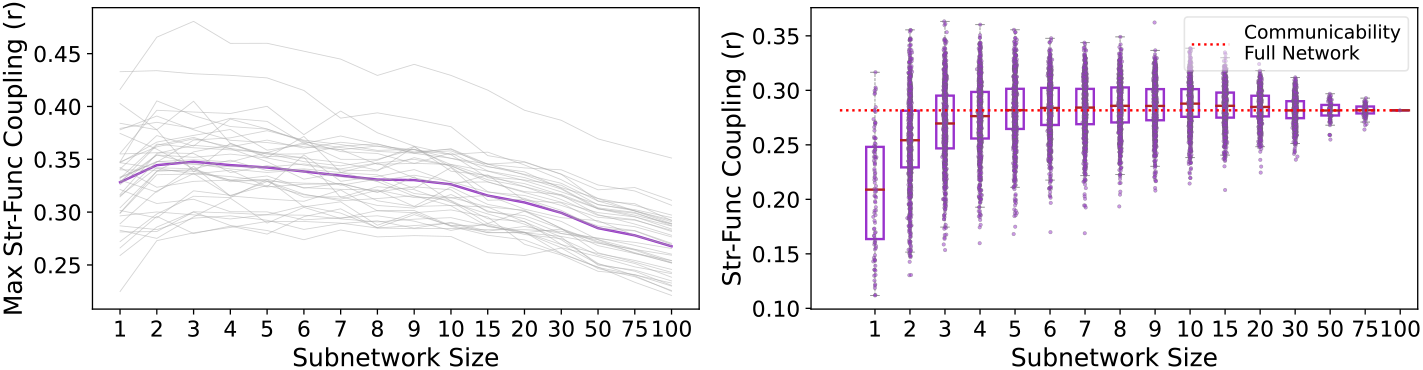
SFC using subnet communicability for varying subnetwork sizes: (left) Highest SFC achieved for each of 40 subjects across varying subnetwork sizes are plotted in gray with their average plotted in purple, demonstrating a peak at subnetworks of size 3 and a decaying SFC with increasing subnetwork sizes. Subnetwork of size 100 corresponds to the putative weighted communicability model **(right)** Distribution of SFC over randomly sampled subnetworks of varying sizes for a single subject demonstrates higher SFC for certain subnetworks relative to standard communicability utilizing the entire network (100 regions).

### Composition of subnetworks

Having empirically determined the subnetwork size with peak SFC, we then analyzed the composition of subnetworks at both the region and system levels. We first investigated brain regions that constitute networks achieving highest SFC. For each of the 200 individuals in the HCP dataset, we randomly sampled 10, 000 subnetworks of size 3 while ensuring connectivity of all regions. We calculated SFC of subnet communicability utilizing each sampled subnetwork as a backbone and evaluated the consistency of subnetwork regions based on how many times a region is included in the subnetwork achieving the highest SFC. As illustrated in Fig. 3, medial prefrontal cortex (MPC) and orbital frontal cortex (OFC) regions were bilaterally over-represented relative to the rest of the regions by several orders of magnitude (Left MPC = 89, right MPC = 98, left OFC = 68, right OFC = 72 occurrences). If region occurrences in a subnetwork were based on random chance, only 6 occurrences would be expected per region across all individuals, thus denoting the significance of the MPC and OFC brain regions. We also noted that only 19 regions were above that threshold.

**Fig. 3.**
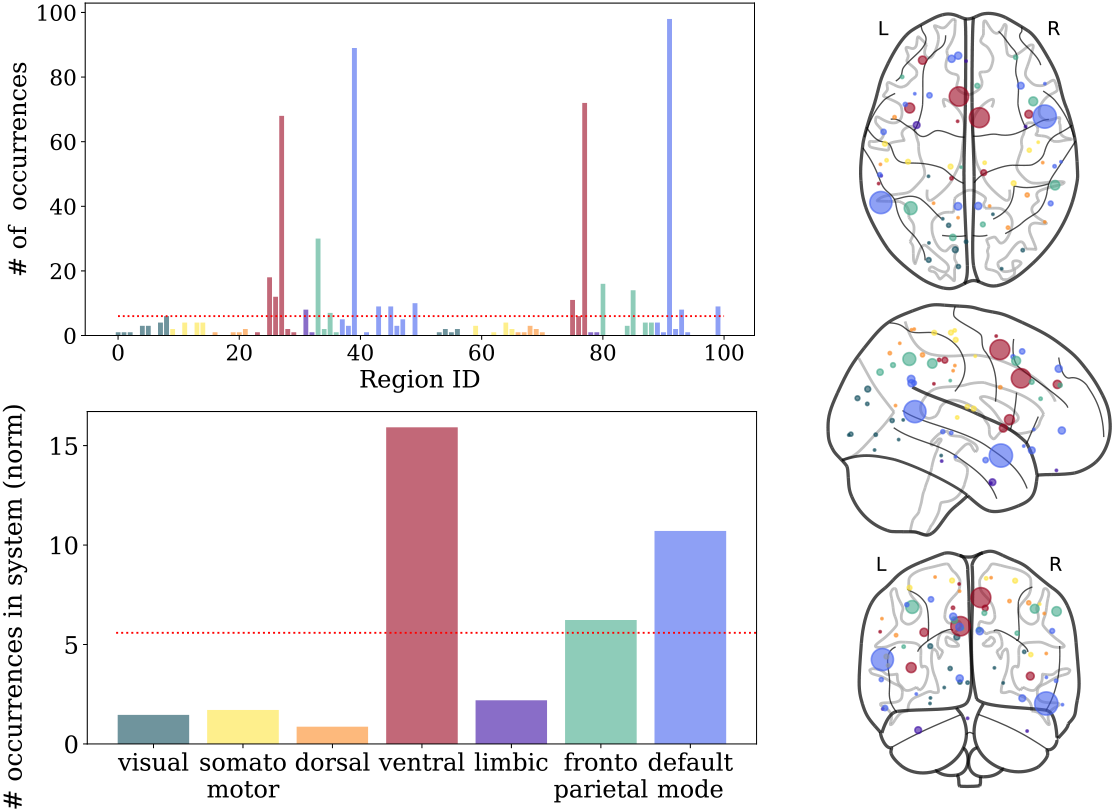
Composition of subnetworks: (top left) Frequency of regions in the highest SFC subnetwork of size 3 across 200 subjects, where *x*-axis denote region IDs. Medial prefrontal cortex and orbital frontal cortex regions bilaterally occurred significantly more than the rest of the regions. **(bottom left)** Frequency of functional systems represented in the highest SFC subnetwork, normalized by system sizes. Default mode and ventral systems are disproportionately overrepresented relative to other systems. (Dashed red lines at the top and bottom indicate the number of times a region is expected to appear in networks across people if the occurrences were by random chance.) **(right)** Occurrences of regions expressed in proportion with the node radius over a brain image.

We then evaluated the representation of the seven functional systems [27] in the highest SFC achieving subnetworks by grouping occurrences of regions into systems followed by a normalization using system size. We observed that the ventral system and default mode network were over-represented while the remaining systems were under-represented (Fig. 3, bottom left). To demonstrate generalizability of results for varying network sizes, we repeated the experiments for subnetworks of size 5 and observed the same pattern at both region and systems levels (Fig. S1 in supplementary material).

### Comparison of communication models

Having established the consistency of certain brain regions in the highest SFC achieving subnetworks, we then explored how subnet communicability performs relative to other communication models. For this subnet communicability analysis, we used the same subnetwork across individuals consisting of bilateral MPC and right OFC regions. We also calculated simulated functional connectomes using the other three communication models. With the putative communicability model as a reference, statistical group comparison (paired t-test) revealed that subnet communicability (*μ* = 0.36) achieved significantly higher SFC (*μ* = 0.27) with a very large effect size (Cohen’s *d* = 2.3, *p <* 10^−6^) (Fig. 4). We further observed that direct communication (*μ* = 0.24; *d* = 0.9, *p <* 10^−6^) and shortest path (*μ* = 0.22; *d* = 1.5, *p <* 10^−6^) both achieve significantly lower SFC, confirming previous literature [17, 20, 28]. Once again, we observed the same pattern of group differences with subnetworks of size 5 (Fig.S2 in supplementary material).

**Fig. 4.**
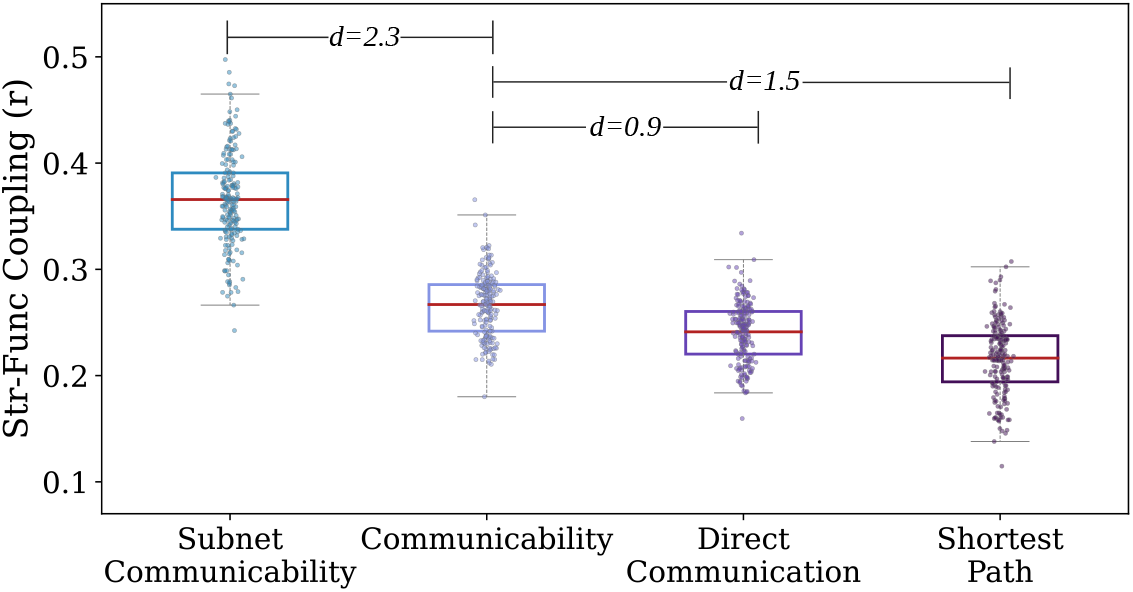
Structure-function coupling of subjects using various communication patterns: Paired group differences were calculated between each communication pattern relative to the putative communicability model. Subnet communicability over a backbone network of 3 nodes achieves significantly higher SFC compared to communicability that uses the entire network for parallel communication. On the other hand, structural connectome without any model applied as well as shortest path achieved a lower SFC relative to communicability. All group differences were significant (*p <* 10^−6^) after Bonferroni multiple comparison correction with large effect sizes (Cohen’s *d*).

### Centrality analysis of subnetworks

Finally, we investigated the network topology features of the highest achieving subnetworks by exploring node centrality measures. We calculated the betweenness (BC), subgraph (SC), eigenvector centralities (EC), and clustering coefficient (CC) of each region over the average SC across subjects. As 19 regions were observed to have representation higher than chance in the composition of subnetworks, we generated 4 sets of 19 regions having the highest network measure scores. We then compared the set of regions that appeared most frequently in subnetworks with these sets to quantify their overlap using the Dice coefficient. We observed low Dice coefficients for all measures (BC=0.32, SC=0.11, EC=0.05, CC=0.11), indicating that region representation in top subnetworks may not be related to the centrality metrics tested alone.

### Replication of results over a second dataset

We repeated the experiments over structural and functional data of 261 individuals from the 1000Brains dataset. The relationship between the size of the subnetwork and SFC demonstrated a trajectory similar to HCP data, where the highest SFC was achieved for networks of size 3-7, and it decreased by larger network sizes (Fig.S3 in supplementary material). When comparing the nodes represented in the subnetwork of size 3 between HCP and 1000Brains datasets, although the individual regions that occurred most frequently were not the same, the nodes from visual, somatomotor, and dorsal systems were under-represented, and ventral and default mode networks were over-represented, which aligns with HCP results. However, we also observed that nodes from limbic and frontoparietal networks were over-represented in this second experiment (Fig.S4 in supplementary material). Finally, repeating the experiment by using the top three most frequently occurring nodes on 1000Brains data, we observed the same pattern of subnet communicability achieving higher SFC than the other communication patterns with large effect sizes (Fig.S5 in supplementary material). Overall, results were replicated to a large extent with some differences in the frequency of occurrences of individual regions.

## 4 Discussion

In this study, we proposed a novel communication model called subnet communicability that uses a considerably smaller subset of brain regions for diffusive message passing. We demonstrated that subnet communicability achieves better structure-function coupling relative to standard communicability over the entire network, which was previously shown to achieve the highest structure-function coupling through a diffusive message passing strategy [17, 28].

The human brain’s wiring is considered to minimize energy consumption [2] supporting communication models such as shortest path. These models deterministically route information using unique optimal paths, requiring global knowledge of network topology, which is deemed unlikely for local brain regions. Diffusive models like communicability, on the other hand, demonstrated higher structure-function coupling while also retaining redundancy in message passing. This approach, thus, ensures robustness in communication while requiring minimal information about network topology. However, communicability contradicts the well-established economy of communication in the brain due to its unrestricted message-passing scheme. Subnet communicability as we proposed, on the other hand, has demonstrated a balance between brain wiring economy and redundancy in message passing while requiring limited knowledge of network topology across neural elements by achieving highest SFC through a fairly small subnetwork of regions.

The over-representation of the default mode network (DMN) and ventral attention system in the subnetworks across subjects demonstrates a consistent pattern. DMN is known to be active during resting state [10] which might explain its central role in subnetworks indicated by the higher SFC relative to resting state function. The ventral attention system is known to orient attention towards internal state [26], which might further support its over-representation for SFC relative to resting state function where external stimuli are limited.

Finally, subnet communicability with the same subnetwork achieving higher SFC consistently across the entire cohort relative to other communication models highlights a biological basis for the choice of nodes that constitute this high-achieving subnetwork. The lack of a strong overlap between the frequency of nodes appearing in highest SFC achieving subnetworks and centrality scores of nodes, however, indicates that tested centrality measures are insufficient in explaining the underlying network topological mechanisms, requiring further investigation.

Although this study investigates the efficacy of subnet communicability over two high-quality datasets, certain limitations should be acknowledged. First, it is known that diffusion MRI has various inaccuracies in determining structural brain connectivity, especially at regions involving crossing or kissing fibers [14]. Second, experiments were carried out on a single atlas at a single parcellation resolution. Third, although the total number of samples investigated in the study is over 500, the datasets explored in the study only consisted of healthy samples, thus limiting the generalizability of results to the healthy. Also, the age range of samples did not include children and young adults, which limits the applicability of results to adults. Finally, prediction of functional interactions of brain regions from their structural connectivity is an inherently restricted problem due to differences of consistency of the data modalities [16].

## 5 Conclusions

Diffusive communication models were previously shown to explain SFC better in the brain and provide necessary redundancy in message passing despite contradicting the established economy of wiring in brain [17, 19, 28]. We have demonstrated that parallel communication when routed through a small subnetwork using subnet communicability explains SFC better than prior diffusive models while establishing a balance between redundancy and economy of brain. Subnet communicability presents an interesting communication pattern that warrants further exploration.

## 6 Acknowledgements

Data was provided by the Human Connectome Project, WU-Minn Consortium (Principal Investigators: David Van Essen and Kamil Ugurbil; 1U54MH091657) funded by the 16 NIH Institutes and Centers that support the NIH Blueprint for Neuroscience Research; and by the McDonnell Center for Systems Neuroscience at Washington University.

## References

1. Avena-Koenigsberger, A., Misic, B., Sporns, O.: Communication dynamics in complex brain networks. Nature reviews neuroscience 19(1), 17–33 (2018)

2. Bullmore, E., Sporns, O.: The economy of brain network organization. Nature reviews neuroscience 13(5), 336–349 (2012)

3. Cabeza, R., Kingstone, A.: Handbook of functional neuroimaging of cognition. Mit Press (2006)

4. Calamante, F., Smith, R.E., Liang, X., Zalesky, A., Connelly, A.: Track-weighted dynamic functional connectivity (tw-dfc): a new method to study time-resolved functional connectivity. Brain Structure and Function 222(8), 3761–3774 (2017)

5. Caspers, S., Moebus, S., Lux, S., Pundt, N., Schütz, H., Mühleisen, T.W., Gras, V., Eickhoff, S.B., Romanzetti, S., Stöcker, T., et al.: Studying variability in human brain aging in a population-based german cohort—rationale and design of 1000brains. Frontiers in aging neuroscience 6, 149 (2014)

6. Craddock, R.C., Jbabdi, S., Yan, C.G., Vogelstein, J.T., Castellanos, F.X., Di Martino, A., Kelly, C., Heberlein, K., Colcombe, S., Milham, M.P.: Imaging human connectomes at the macroscale. Nature methods 10(6), 524–539 (2013)

7. Crofts, J.J., Higham, D.J.: A weighted communicability measure applied to complex brain networks. Journal of the Royal Society Interface 6(33), 411–414 (2009)

8. Domhof, J.W.M., Jung, K., Eickhoff, S.B., Popovych, O.V.: Parcellation-based structural and resting-state functional brain connectomes of a healthy cohort (v1.1) (2022). 10.25493/NVS8-XS5, https://search.kg.ebrains.eu/instances/f16e449d-86e1-408b-9487-aa9d72e39901

9. Estrada, E., Hatano, N.: Communicability in complex networks. Physical Review E 77(3), 036111 (2008)

10. Fox, M.D., Snyder, A.Z., Vincent, J.L., Corbetta, M., Van Essen, D.C., Raichle, M.E.: The human brain is intrinsically organized into dynamic, anticorrelated functional networks. Proceedings of the National Academy of Sciences 102(27), 9673–9678 (2005)

11. Goñi, J., Van Den Heuvel, M.P., Avena-Koenigsberger, A., Velez de Mendizabal, N., Betzel, R.F., Griffa, A., Hagmann, P., Corominas-Murtra, B., Thiran, J.P., Sporns, O.: Resting-brain functional connectivity predicted by analytic measures of network communication. Proceedings of the National Academy of Sciences 111(2), 833–838 (2014)

12. Honey, C.J., Sporns, O., Cammoun, L., Gigandet, X., Thiran, J.P., Meuli, R., Hagmann, P.: Predicting human resting-state functional connectivity from structural connectivity. Proceedings of the National Academy of Sciences 106(6), 2035–2040 (2009)

13. Honey, C.J., Thivierge, J.P., Sporns, O.: Can structure predict function in the human brain? Neuroimage 52(3), 766–776 (2010)

14. Jones, D.K.: Challenges and limitations of quantifying brain connectivity in vivo with diffusion mri. Imaging in Medicine 2(3), 341 (2010)

15. Jung, K., Eickhoff, S.B., Popovych, O.V.: Parcellation-based structural and resting-state functional whole-brain connectomes of 1000brains cohort (v1.1) (2022). 10.25493/8XY5-BH7, https://search.kg.ebrains.eu/instances/3f179784-194d-4795-9d8d-301b524ca00a

16. Osmanlıoğlu, Y., Alappatt, J.A., Parker, D., Verma, R.: Connectomic consistency: a systematic stability analysis of structural and functional connectivity. Journal of neural engineering 17(4), 045004 (2020)

17. Osmanlıoğlu, Y., Tunç, B., Parker, D., Elliott, M.A., Baum, G.L., Ciric, R., Satterthwaite, T.D., Gur, R.E., Gur, R.C., Verma, R.: System-level matching of structural and functional connectomes in the human brain. NeuroImage 199, 93–104 (2019)

18. Schaefer, A., Kong, R., Gordon, E.M., Laumann, T.O., Zuo, X.N., Holmes, A.J., Eickhoff, S.B., Yeo, B.T.: Local-global parcellation of the human cerebral cortex from intrinsic functional connectivity mri. Cerebral cortex 28(9), 3095–3114 (2018)

19. Seguin, C., Jedynak, M., David, O., Mansour L, S., Sporns, O., Zalesky, A.: Communication dynamics in the human connectome shape the cortex-wide propagation of direct electrical stimulation. bioRxiv pp. 2022–07 (2022)

20. Seguin, C., Sporns, O., Zalesky, A., Calamante, F., et al.: Network communication models narrow the gap between the modular organization of structural and functional brain networks. NeuroImage 257, 119323 (2022)

21. Seguin, C., Tian, Y., Zalesky, A.: Network communication models improve the behavioral and functional predictive utility of the human structural connectome. Network Neuroscience 4(4), 980–1006 (2020)

22. Seguin, C., Van Den Heuvel, M.P., Zalesky, A.: Navigation of brain networks. Proceedings of the National Academy of Sciences 115(24), 6297–6302 (2018)

23. Sporns, O., Tononi, G., Kötter, R.: The human connectome: a structural description of the human brain. PLoS computational biology 1(4), e42 (2005)

24. Van Den Heuvel, M.P., Kahn, R.S., Goñi, J., Sporns, O.: High-cost, high-capacity backbone for global brain communication. Proceedings of the National Academy of Sciences 109(28), 11372–11377 (2012)

25. Van Essen, D.C., Ugurbil, K., Auerbach, E., Barch, D., Behrens, T.E., Bucholz, R., Chang, A., Chen, L., Corbetta, M., Curtiss, S.W., et al.: The human connectome project: a data acquisition perspective. Neuroimage 62(4), 2222–2231 (2012)

26. Vossel, S., Geng, J.J., Fink, G.R.: Dorsal and ventral attention systems: distinct neural circuits but collaborative roles. The Neuroscientist 20(2), 150–159 (2014)

27. Yeo, B.T., Krienen, F.M., Sepulcre, J., Sabuncu, M.R., Lashkari, D., Hollinshead, M., Roffman, J.L., Smoller, J.W., Zöllei, L., Polimeni, J.R., et al.: The organization of the human cerebral cortex estimated by intrinsic functional connectivity. Journal of neurophysiology (2011)

28. Zamani Esfahlani, F., Faskowitz, J., Slack, J., Mišić, B., Betzel, R.F.: Local structure-function relationships in human brain networks across the lifespan. Nature communications 13(1), 1–16 (2022)

