## Supplementary Information for "Subnet Communicability: Diffusive Communication Across the Brain Through a Backbone Subnetwork"

### 1 Repeating experiments over a subnetwork consisting of five nodes on HCP data

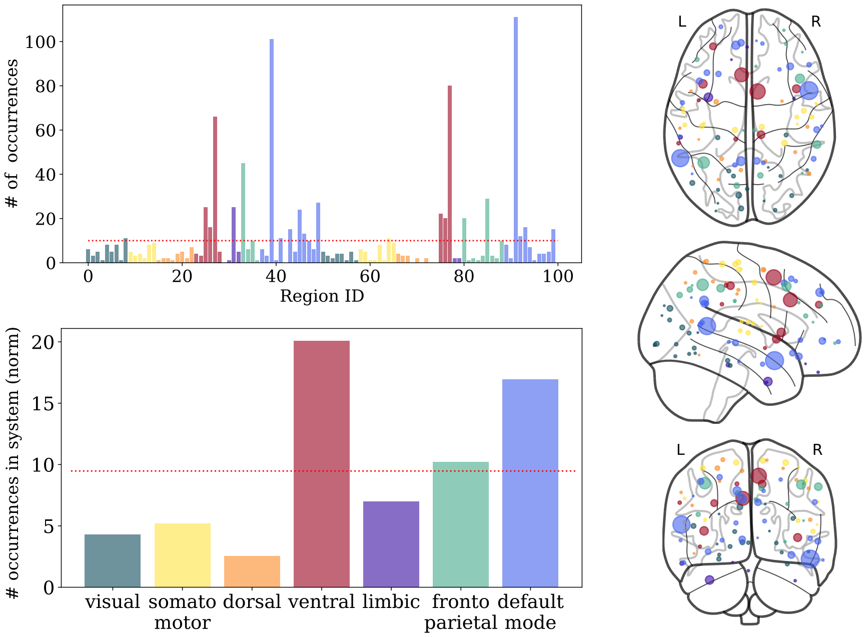


**Fig. 1. Composition of subnetworks: (top left)** Frequency of regions in the highest SFC subnetwork of size 5 across 200 subjects, where $x$-axis denote region IDs. Bilateral medial prefrontal cortices and orbital frontal cortices, and left lateral ventral prefrontal cortex regions occurred significantly more than the rest of the regions. **(bottom left)** Frequency of functional systems represented in the highest SFC subnetwork, normalized by system sizes. Default mode and ventral systems are disproportionately over represented relative to other systems. (Dashed red lines at top and bottom indicates the number of times a region is expected to appear in networks across people if the occurrences were by random chance.) **(right)** Occurrences of regions expressed in proportion with the node radius over a brain image.

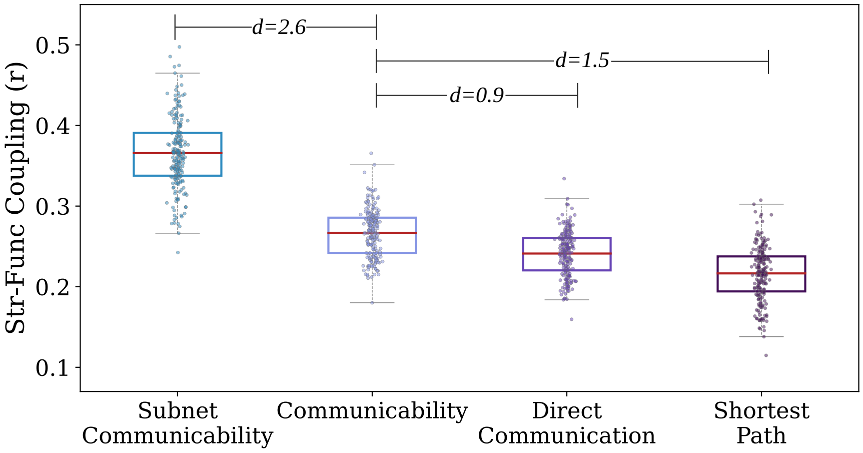


**Fig. 2. Structure-function coupling of subjects using various communication patterns:** Paired group differences were calculated between each communication pattern relative to the putative communicability model. Subnet communicability over a backbone network of 5 nodes achieves significantly higher SFC compared to communicability that uses the entire network for parallel communication. On the other hand, structural connectome without any model applied as well as shortest path achieved a lower SFC relative to communicability. All group differences were significant ($p < 10^{-6}$) after Bonferroni multiple comparison correction with large effect sizes (Cohen's $d$).

### 2 Replication Study: Repeating experiments over a subnetwork consisting of three nodes on 1000Brains dataset

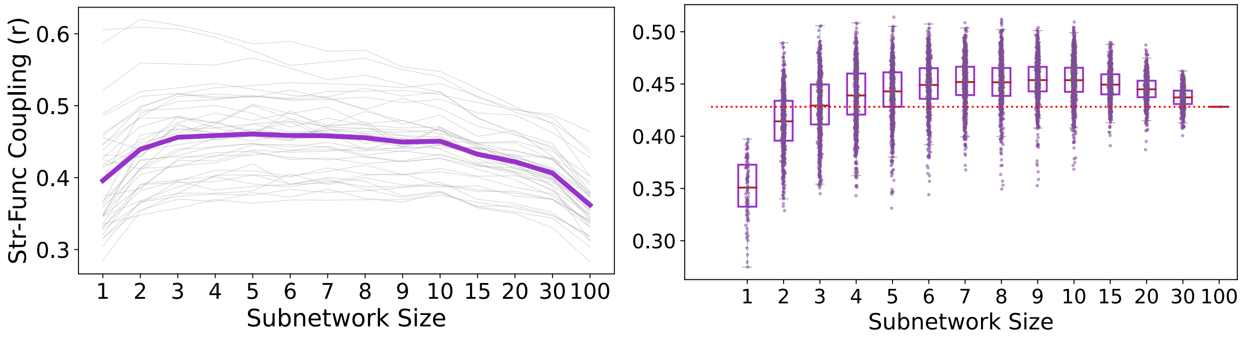


**Fig. 3. SFC using subnet communicability for varying subnetwork sizes over 1000Brains dataset: (left)** Highest SFC achieved for each of 40 subjects across varying subnetwork sizes are plotted in gray with their average plotted in purple, demonstrating a peak at subnetworks of size 3-7 and a decaying SFC with increasing subnetwork sizes. Subnetwork of size 100 corresponds to the putative weighted communicability model **(right)** Distribution of SFC over randomly sampled subnetworks of varying sizes for a single subject demonstrates higher SFC for certain subnetworks relative to standard communicability utilizing the entire network (100 regions).

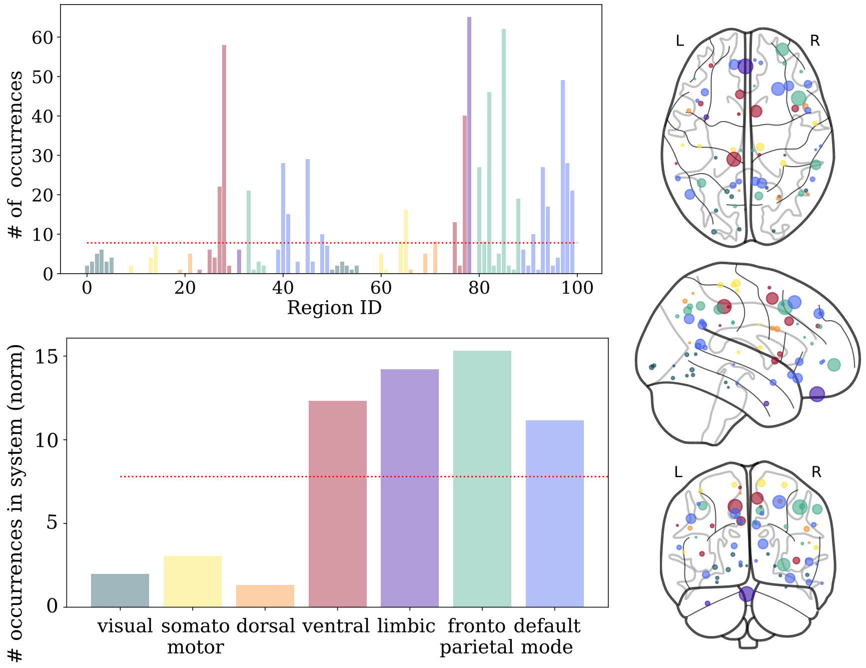


**Fig. 4. Composition of subnetworks over 1000Brains dataset: (top left)** Frequency of regions in the highest SFC subnetwork of size 3 across 261 subjects, where $x$-axis denote region IDs. Salient Ventral Attention Med-2 region on left hemisphere and Limbic Orbitofrontal Cortext-1 and PFCl-4 regions on the right hemispheres are top three most frequently chosen regions to constitute the subnetwork. **(bottom left)** Frequency of functional systems represented in the highest SFC subnetwork, normalized by system sizes. Limbic, frontoparietal, default mode, and ventral systems are disproportionately over represented relative to other systems. (Dashed red lines at top and bottom indicates the number of times a region is expected to appear in networks across people if the occurrences were by random chance.) **(right)** Occurrences of regions expressed in proportion with the node radius over a brain image.

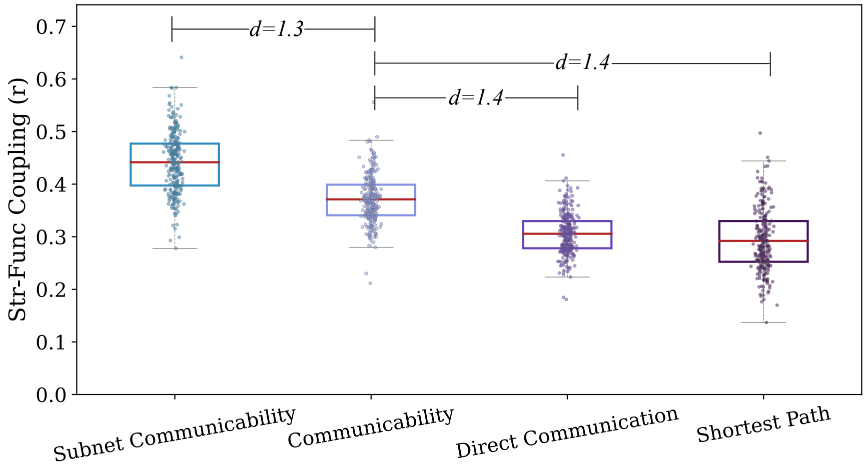


**Fig. 5. Structure-function coupling of subjects using various communication patterns on 1000Brains dataset:** Paired group differences were calculated between each communication pattern relative to the putative communicability model. Subnet communicability over a backbone network of three nodes achieves significantly higher SFC compared to communicability that uses the entire network for parallel communication. On the other hand, structural connectome without any model applied as well as shortest path achieved a lower SFC relative to communicability. All group differences were significant ($p < 10^{-6}$) after Bonferroni multiple comparison correction with large effect sizes (Cohen's $d$).
